# Compartmentalized three-dimensional human neuromuscular tissue models fabricated on a well-plate-format microdevice

**DOI:** 10.1101/2021.01.07.424253

**Authors:** Kazuki Yamamoto, Nao Yamaoka, Yu Imaizumi, Takunori Nagashima, Taiki Furutani, Takuji Ito, Yohei Okada, Hiroyuki Honda, Kazunori Shimizu

## Abstract

Engineered three-dimensional models of neuromuscular tissues are promising for use in mimicking their disorder states in vitro. Although several models have been developed, it is still challenging to mimic the physically separated structures of motor neurons (MNs) and skeletal muscle (SkM) fibers in the motor units in vivo. In this study, we aimed to develop microdevices for precisely compartmentalized coculturing of MNs and engineered SkM tissues. The developed microdevices, which fit a well of 24 well plates, had a chamber for MNs and chamber for SkM tissues. The two chambers were connected by microtunnels for axons, permissive to axons but not to cell bodies. Human iPSC (hiPSC)-derived MN spheroids in one chamber elongated their axons into microtunnels, which reached the tissue-engineered human SkM in the SkM chamber, and formed functional neuromuscular junctions with the muscle fibers. The cocultured SkM tissues with MNs on the device contracted spontaneously in response to spontaneous firing of MNs. The addition of a neurotransmitter, glutamate, into the MN chamber induced contraction of the cocultured SkM tissues. Selective addition of tetrodotoxin or vecuronium bromide into either chamber induced SkM tissue relaxation, which could be explained by the inhibitory mechanisms. We also demonstrated the application of chemical or mechanical stimuli to the middle of the axons of cocultured tissues on the device. Thus, compartmentalized neuromuscular tissue models fabricated on the device could be used for phenotypic screening to evaluate the cellular type specific efficacy of drug candidates and would be a useful tool in fundamental research and drug development for neuromuscular disorders.

## 1. Introduction

The physical movement of the body is induced by the transmission of signals to the motor units, consisting of motor neurons (MNs) and skeletal muscle (SkM) fibers. MNs whose cell body is in the spinal cord elongate the axon to the SkM fibers. The terminals of axons form neuromuscular junctions (NMJs), chemical synapses, with muscle fibers. When an action potential reaches the terminal of the axon, acetylcholine is released from the axon terminal and binds to the acetylcholine receptors (AChRs) on the muscle fiber membrane, inducing muscle contraction (1, 2). Neuromuscular disorders such as MN diseases, neuropathies, NMJ disorders, and myopathies affect one or more components of the motor unit, causing progressive motility disturbances (3). As neuromuscular disorders span a broad range of rare conditions with diverse genetic and nongenetic etiologies and pathophysiologies, it is particularly challenging to develop comprehensive animal models (4). Thus, it is required to develop more efficient and predictive *in vitro* neuromuscular disorder models to improve our mechanistic knowledge on these disorders and develop effective treatments (4–7).

Engineered three-dimensional (3D) models of neuromuscular tissues are promising for use in mimicking the disorder state *in vitro*. A common approach to fabricate them is to attach/mix the MNs directly to the tissue-engineered SkM (8–13). The usefulness of this approach for modeling of neuromuscular disorders has been demonstrated by showing that the addition of IgG isolated from myasthenia gravis patients, an autoimmune disease of NMJs, reduced the contraction of the innervated tissues (13). However, owing to the attaching/mixing nature of this approach, it is challenging to monitor individual cell types and control their culture conditions.

To overcome this issue, another approach has been proposed where the coculture of MNs and engineered SkM tissues with their cell bodies is physically separated, which mimics the physically separated structure of MNs and muscle fibers in the motor units (14–17). In the pioneering studies by Kamm et al., microfluidic devices for the formation of compartmentalized motor units were developed (14, 15). MN spheroids and engineered SkM tissues were positioned separately in a microchannel filled with collagen gel. The axons of MNs elongated within the gel reached the muscle tissues and formed NMJs with the muscle fibers in the tissues. The devices have been used to model amyotrophic lateral sclerosis using induced pluripotent stem cells (iPSCs) from patients. This indicates that this approach has the potential for mimicking neuromuscular disorders. However, using the reported devices, it is still challenging to keep all cell bodies of MNs and muscle fibers separated during a culture period and precisely control the culture conditions for each cell type because of the rough compartmentalization by the collagen gel.

Compartmentalized coculture of cell bodies and axons of neurons has been achieved by microfabricated devices with two chambers connected to each other by microtunnels permissive to axons but not to cell bodies (18, 19). Devices with microtunnels have been applied in development of *in vitro* two-dimensional (2D) compartmentalized motor unit models by coculturing MNs and SkM cells in each chamber (20–23). The isolation characteristic of the chambers allows to add drugs or other small molecules to one or both chambers selectively (24). Electrical stimulation can be applied to the cells in each chamber (25). Furthermore, it is possible to apply chemical and/or physical stimulation to the middle of the axons to study axon injury and regeneration (26). Thus, compartmentalization by the microtunnels for axons has been widely accepted as a powerful approach for neuromuscular research.

In this study, we propose microdevices for a precisely compartmentalized coculture of MNs and engineered SkM tissues (Fig. 1). We integrated the microtunnels for axons into the microdevices for a contractile force measurement of the developed engineered SkM tissues (27–29). The developed microdevices that fit a well of a 24-well plate have two chambers connected by the microtunnels for axons. Human iPSC (hiPSC)-derived MN spheroids in one chamber elongated their axons into microtunnels, which reached the tissue-engineered human SkM in the other chamber, and formed functional NMJs with the muscle fibers. As the cell bodies of the MNs and SkM cells were separated by the microtunnels, we succeeded in controlling the culture conditions of each cell type and applied chemical or physical stimulations to cell bodies or middle of the axons selectively. To the best of our knowledge, this is the first study on microdevices for the fabrication of precisely compartmentalized 3D models of neuromuscular tissues.

**Fig. 1.**
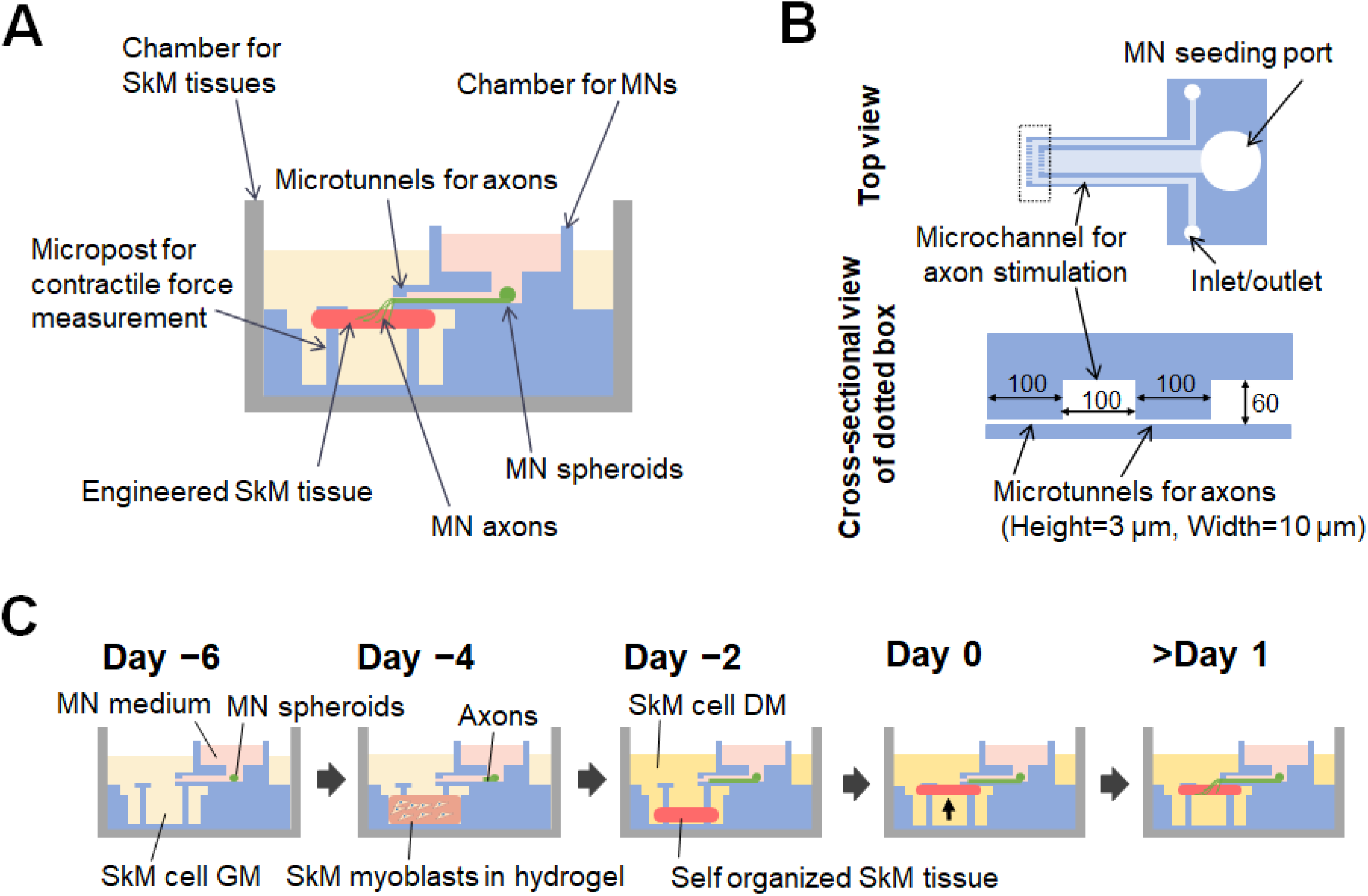
Schematic illustration of the microdevice. A) Cross sectional view. B) A chamber for MNs. C) Scheme for coculture on the microdevices.

## 2. Materials and Methods

### 2.1 Materials

SU-8 3005 (Nippon Kayaku, Tokyo, Japan), SU-8 developer (Nippon Kayaku, Tokyo, Japan), polydimethylsiloxane (PDMS; SILPOT 184, DuPont Toray Specialty Materials K.K., Tokyo, Japan), StemFit (AK02N, Ajinomoto, Tokyo, Japan), i-Matrix-511 (381-07363, Fujifilm Wako Chemicals, Osaka, Japan), TrypLE™ Select CTS™ (A12859-01, Thermo Fisher Scientific, Waltham, MA), UltraPure™ 0.5-M EDTA (15575020, Thermo Fisher Scientific), Y-27632 (08945-71, Nacalai Tesque, Kyoto, Japan), fibrinogen (F8630, Sigma-Aldrich), thrombin from bovine plasma (T4648, Sigma-Aldrich, St. Louis, MO), 6-aminocapuroic acid (6-AA, A2504, Sigma-Aldrich), trans-4-(aminomethyl) cyclohexanecarboxylic acid (TA, A0236, Tokyo Chemical Industry, Tokyo, Japan), DMEM (08458-16, Nacalai tesque), Ultroser G serum substitute (15950-017, Pall Corporation, Port Washington, NY, USA), Matrigel® basement membrane matrix (354234, Corning Inc., Corning, NY, USA), Dulbecco’s modified Eagle medium/ Nutrient Mixture F-12 (DMEM-F12; 042-30555, Fujifilm Wako Chemicals), pluronic F-127 (low ultraviolet (UV) absorbance) (P6867, Thermo Fisher Scientific), tetrodotoxin (TTX, 206-11071, Fujifilm Wako Chemicals), vecuronium bromide (Vb; 223-01811, Fujifilm Wako Chemicals) were employed.

### 2.2 Microdevice design and fabrication

The microdevice consisted of a chamber for SkM tissues and chamber for MNs (Fig. 1). The SkM tissue chamber fabricated in our previous study (27) and MN chamber developed in this study were integrated. The MN chamber was fabricated by standard PDMS soft lithography using an SU-8 mold. Micropatterned molds were fabricated by negative photoresist photolithography using SU-8 on a silicon wafer. To fabricate microtunnels for axons with a height of 3 μm, SU-8 3005 was diluted (1:1) with cyclopentanone, spin-coated onto a Si wafer at 3000 rpm for 30 s, and heated at 95 °C for 10 min. The wafer was then exposed to UV light through a photomask and heated at 95 °C for 10 min. Subsequently, to fabricate microchannels with a height of 60 μm, SU-8 3050 was spin-coated on the silicon wafer with micropatterns at 2000 rpm for 30 s, heated at 95 °C for 1 h, exposed to UV light through a second mask, and then heated at 95 °C for 10 min. The wafers were developed in an SU-8 developer and washed with isopropanol. PDMS (base:catalyst = 10:1) was placed on the mold, spin-coated at 800 rpm for 30 s, and heated at 70 °C for 1 h to fabricate part I (Supplementary Fig. 1). A cylindrical PDMS (diameter: 8 mm, height: 2 mm) (Part II) was attached to Part I by an air plasma treatment, and then the MN seeding port and inlet/outlet for the microchannel for axon stimulation were punched out with biopsy punches with diameters of 2 and 1.5 mm, respectively. After cutting the tip of part I, a part-III cylindrical PDMS (outer diameter: 8 mm, height: 2 mm) with a hole (diameter: 6 mm) was bonded to part II. The assembled parts were then bonded to the PDMS film (thickness of approximately 30 μm) on Si wafers using air plasma. The fabricated MN chamber was bonded onto the chamber for SkM tissues (27) The completed microdevices were exposed to UV light for at least 2 h for sterilization before use for the coculture.

### 2.3 Cell culture

hiPSC clone (201B7) (30) were provided by Dr. Shinya Yamanaka (RIKEN BRC through the Project for Realization of Regenerative Medicine and National Bio-Resource Project of the MEXT, Japan), and cultured as reported previously (31). hiPSCs were maintained with Stemfit AK02N on an iMatrix-511 coating (0.5 μg/cm^2^). MN differentiation of hiPSCs was performed according to a reported method (32). hiPSCs were stripped from the flask with a 1:1 mixture of TrypLE:0.5-mM EDTA. The cells were suspended at 1.0 × 10^5^ cells/mL in AK02N containing 10-μM Y27632 and seeded into a 6-mm cell culture dish to form spheroids. After the spheroid was formed, the medium was exchanged with embryoid body medium and cultured for 14 days before seeding to the device. Hu5/KD3 is an immortalized human myogenic cell developed by Hashimoto et al. (33). The Hu5/KD3 myoblasts were seeded into collagen-type-I-coated T-75 flasks and cultured at 37 °C under 5% CO_2_ and 10% O_2_ in DMEM consisting of 20% FBS, 0.5% penicillin–streptomycin, 2 mM of L-glutamine, and 2% Ultroser G serum substitute (GM).

### 2.4 3D coculture of SkM tissues and MN spheroids on the microdevices

The process for the coculture is shown in Fig. 1C. To prevent adhesion of muscle tissue to the surface of the microdevice, the dumbbell-shaped pockets of the device were filled with a 2% Pluronic solution and incubated for 1 h at 25 °C. After coating, the solution was removed and the entire device was soaked in a Matrigel solution diluted 30-fold by DMEM-F12 and incubated at 25 °C for 3 h to enable MN adhesion onto the surface of the MN chamber. After all solutions were drained, hiPSC-derived MN spheroids were seeded into the MN seeding port on day −6. The chamber for MNs was filled with a MN medium, while the SkM tissue chamber was filled with GM. On day −4, an SkM tissue was fabricated, according to the reported method (29). A hydrogel containing fibrin, Matrigel, and human SkM myoblasts, Hu5/KD3 myoblasts, were formed by adding 2% thrombin at a volume of 1.6% to the solution, filling a dumbbell-shaped pocket, and incubating at 37 °C for 30 min. GM was replaced with GM with 6-AA and TA to prevent hydrogel degradation. On day −2, the medium was replaced with DMEM containing 2% horse serum, 1% PS, 2 mg of 6-AA, and 1-mg/mL TA (differentiation medium (DM)). The medium was replaced every two or three days. On day 0, the SkM tissues were lifted up to the top of the microposts for contact with the tip of the MN chamber. Subsequently, we observed contraction of the SkM tissues for 10 days.

### 2.5 Evaluation of the contractility of SkM tissues cocultured with MNs

The displacement of the tip of the micropost was observed under an upright microscope (BX53, OLYMPUS, Japan) equipped with ORCA-Spark (Hamamatsu Photonics, Hamamatsu, Japan). The contractile force was calculated using the displacement of the tip using the reported equation (29). For the measurement of contraction of the cocultured SkM tissues induced by spontaneous firing of MNs, the displacement of the tip of the micropost was observed within 5 min after removing the tissues from the 37 °C incubator to avoid the loss of MN activities due to the temperature decrease. To activate the MNs, glutamate was added at 400 μM into the MN chamber. TTX or Vb was added at 2 μM or 20 μM, respectively, into either the SkM chamber or MN chamber, immediately after the addition of glutamate into the MN chamber. A motion tracking software (PV Studio 2D, OA Science, Miyazaki, Japan) was used to analyze the movement of the tip of the micropost.

### 2.6 Region-selective axonal stimulation on the microdevice

The staining solution was prepared by adding 5 μL of Cell Brite™ Membrane Dyes to 1 mL of culture medium. The medium was removed from the microdevice and 7 μL of the staining solution was slowly injected into the inlet of the microchannel for axon stimulation (Fig. 1B). After incubation at 37 °C for 30 min, the microchannel was washed with PBS and observed under a fluorescence microscope (BZ-X700, Keyence).

To denature the axons, shear stress was applied by flowing liquid through the microchannel for axon stimulation. The tip of the microchip was placed in a close contact with the inlet/outlet of the microfluidic channel and the culture medium was forced to flow (Fig. 1B). The shear stress was calculated as

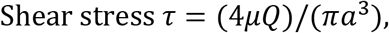

where *μ* is the viscosity of the flowing liquid [Pa·s], *Q* is the volume flow rate [m^3^/s], and *a* is the radius of the channel [m]. As the microchannel is rectangular, the equivalent radius was obtained by dividing the cross-sectional area by the perimeter.

## 3. Results and Discussion

### 3.1 Concept and design of the microdevices

A conceptual illustration of the developed microdevice is shown in Fig. 1. The microdevice had two chambers, one for engineered SkM tissues and other for MNs (Fig. 1A). As the chambers are separated by the microtunnels for axons, the axons of MNs seeded into the MN chamber reach the SkM tissues cultured in the SkM chamber. The contractile force of the SkM tissues is quantified by measuring the displacement of the tip of the micropost in response to muscle contraction (29). Two steps of microtunnels for axons were designed at the tip of the MN chamber (Fig. 1B). The height and width of the microtunnels for the axons were designed to be 3 and 10 μm, respectively. To selectively apply chemical or mechanical stimuli to the middle of the axons, the MN chamber had a microchannel for axon stimulation (Fig. 1B).

The developed device is shown in Fig. 2A. The device was compatible with the well of a commercially available 24-well plate (Fig. 2B). The tips of the MN chambers are shown in Fig. 2C. We developed a MN chamber with two steps of three and nine microtunnels for the axons. The axons elongating in the MN chamber (+) pass through the three and nine microchannels for the axon simultaneously and reach the outside space (*) of the MN chamber. The gap between the microtunnels for axons (#) acts as a microchannel for axon stimulation. Fig. 2D and E show representative images of the coculture on the microdevice in a well of the 24-well plate. The ribbon-shaped innervated SkM tissue was formed by self-organization.

**Fig. 2.**
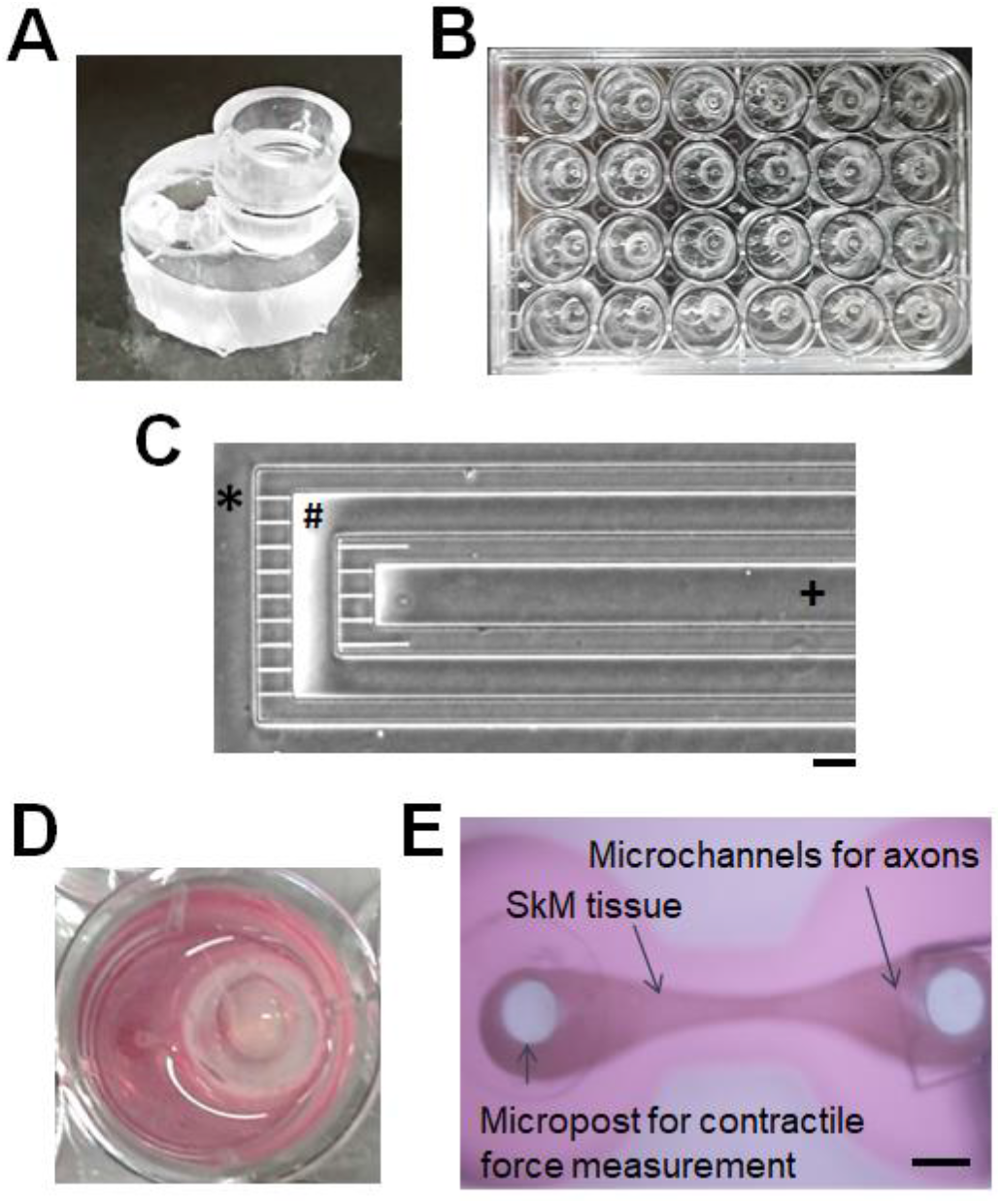
Fabrication and use of the microdevice. A) Image of the fabricated microdevice. B) Image of the 24 well plate with the microdevices. C) Top view image of the MN chamber. Axons from MN spheroid elongate in the MN chamber (+) and reached to the outside space (*). The gap between the microtunnels for axons (#) is used as a microchannel for axon stimulation. Scale bar; 100 μm. D) Image of a well of 24 well plate with the device with SkM tissues and MNs. E) Image of the SkM tissue on the device. Scale bar; 500 μm.

### 3.2 Cell culture using MN chambers

We investigated whether compartmentalized culturing was possible using the developed MN chamber. hiPSC-derived MN spheroids were seeded in the MN seeding port in the MN chamber. After six days of culturing, the MN elongated the axons in the microchannel (+) and the tips of several axons reached the three microtunnels for axons (Fig. 3A). After 10 days, the tips of dozens of axons passed through the nine microtunnels for axons and reached the outside (*) of the MN chamber (Fig. 3B). Notably, the axons in the MN chamber (+) formed a fork-like structure with three tines by self-assembling of axons (Fig. 3B). We confirmed that the number of tines was consistent with the number of microtunnels using the MN chamber with one step of nine microtunnels for the axon (Supplementary Fig. 2). Notably, no cell bodies of MNs were observed in the areas of (#) and (*) (Fig. 3B). Furthermore, we confirmed that the SkM myoblasts cultured in the area (*) did not migrate into the area (#). These results suggest that compartmentalized coculturing of MNs and SkM tissues could be achieved using the developed MN chamber.

**Fig. 3.**
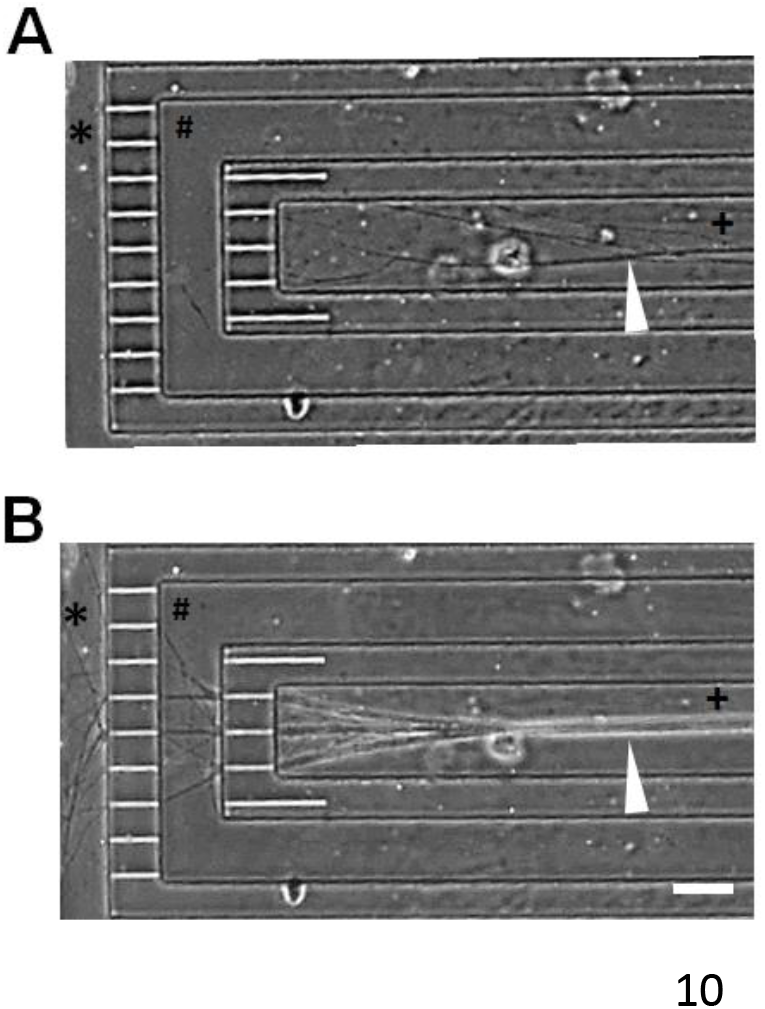
Axon elongation in the MN chamber. Representative image of the elongated axons from MN spheroid on A) day6 and B) day 10. Anow head; elongated axons. Axons from MN spheroid elongate in the MN chamber (+) and reached to the outside space (*). The gap between the microtmmels for axons (#) is used as a microchannel for axon stimulation. Scale bar; 100 μm.

The structure of the fork handle seemed to be similar to the structure of the axon fascicle, as reported by Kawada et al. (34). They reported that the axons extended from the iPSC-derived MN spheroid in the Matrigel-coated PDMS microchannel (width = 150 μm and height = 150 μm) spontaneously assembled into a unidirectional fascicle; the axon fascicle was electrically active and elastic. This suggests that the fork-like self-assembled axons are also electrically active and that the MN chamber could be used to culture functional MN spheroids with an axon fascicle.

### 3.3 Selective stimulation of the middle of the axons of the MNs cultured in the MN chamber

Further, we evaluated whether chemical or mechanical stimulation could be applied selectively to the axons of the MNs elongated in the microchannel for axon stimulation of the MN chamber. MN spheroids were seeded into the MN seeding port and cultured until their axons reached the outside of the MN chamber. For the chemical stimulation, 7 μL of the medium mixed with Cell Brite^TM^ cytoplasmic membrane dye was injected slowly into the microchannel for axon stimulation (#) (Fig. 4A). After 30 min, the fluorescently stained axons were observed in the area (#), whereas the axons in the area (+) were not stained (Fig. 4B). These results suggest that selective chemical stimulation was possible. For the mechanical stimulation, 200 μL of the medium was injected at a shear stress of 60 Pa into the microchannel for axon stimulation (#) (Fig. 4C). The elongated axons in the area (#) disappeared after the application of the shear stress for 30 s, whereas the axons in the area (+) remained as they were (Fig. 4D). Notably, the axons reelongated in the area (#) after 24 h of stimulation, which suggests a possible application of the developed coculture microdevice to model the axon injury and regeneration (Fig. 4E). These results indicate that mechanical stimuli can be applied selectively to the axons of the MN in the microchannel for axon stimulation in the MN chamber.

**Fig. 4.**
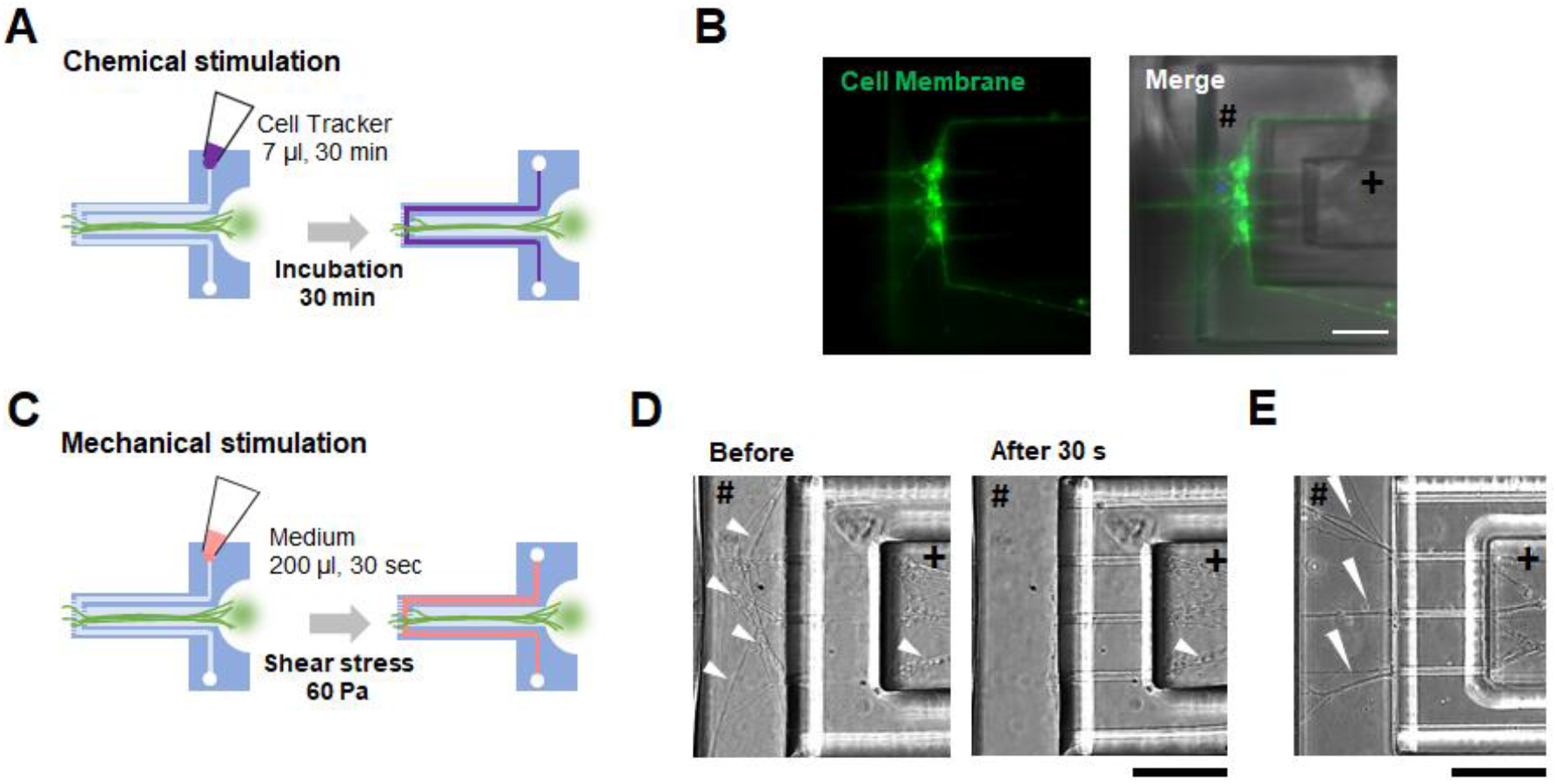
Selective stimulation of the middle of the axons. A) Scheme for chemical stimulation. B) Selective fluorescent labeled axons in the microchannel for axon stimulation (#). Scale bar; 100 μm. C) Scheme for mechanical stimulation. D) Selective removal of the axons in the microchannel for axon stimulation (#) by shear stress. Scale bar; 100 μm. E) Re-elongation of the axons in the microchannels for axon stimulation (#). Scale bar; 100 μm.

### 3.4 Time course trend of spontaneous contraction of engineered SkM tissues cocultured with MN spheroids on the device

We cocultured MN spheroids and engineered SkM tissues on the device. SkM myotubes innervated with MNs exhibited spontaneous contraction in response to spontaneous firing of MNs (35). Thus, we analyzed the spontaneous contraction of the SkM tissues cocultured with MN spheroids on the devices.

Fig. 5A shows the appearance of the cocultured SkM tissues on the device on days 0, 4, and 8. The tip of the micropost was gradually pulled toward the MN chamber by increasing the passive tension (Fig. 5B). The passive tension of the cocultured SkM tissue was significantly larger than that of the monocultured SkM tissue. Fig. 5C shows the displacement of the tip of the micropost moved by the spontaneous contraction of the SkM tissue on days 4, 6, and 8. The SkM tissue contracted periodically at approximately 2 Hz on days 4 and 6 and nonperiodically on day 8. Notably, the displacements were largely different between the days. Thus, the contractile force was calculated using the displacement. The time course trend of the contractile force is shown in Fig. 5D. The contractile force was 0.92 ± 0.18 μN on day 2 and slightly increased to 1.9 ± 0.052 μN on day 4. It largely increased to 18 ± 4.2 μN on day 5 and was maintained at 19 ± 1.7 μN on day 6. It then decreased to 0.25 ± 0.028 μN on day 8 and remained at 0.25 ± 0.023 μN on day 10.

**Fig. 5.**
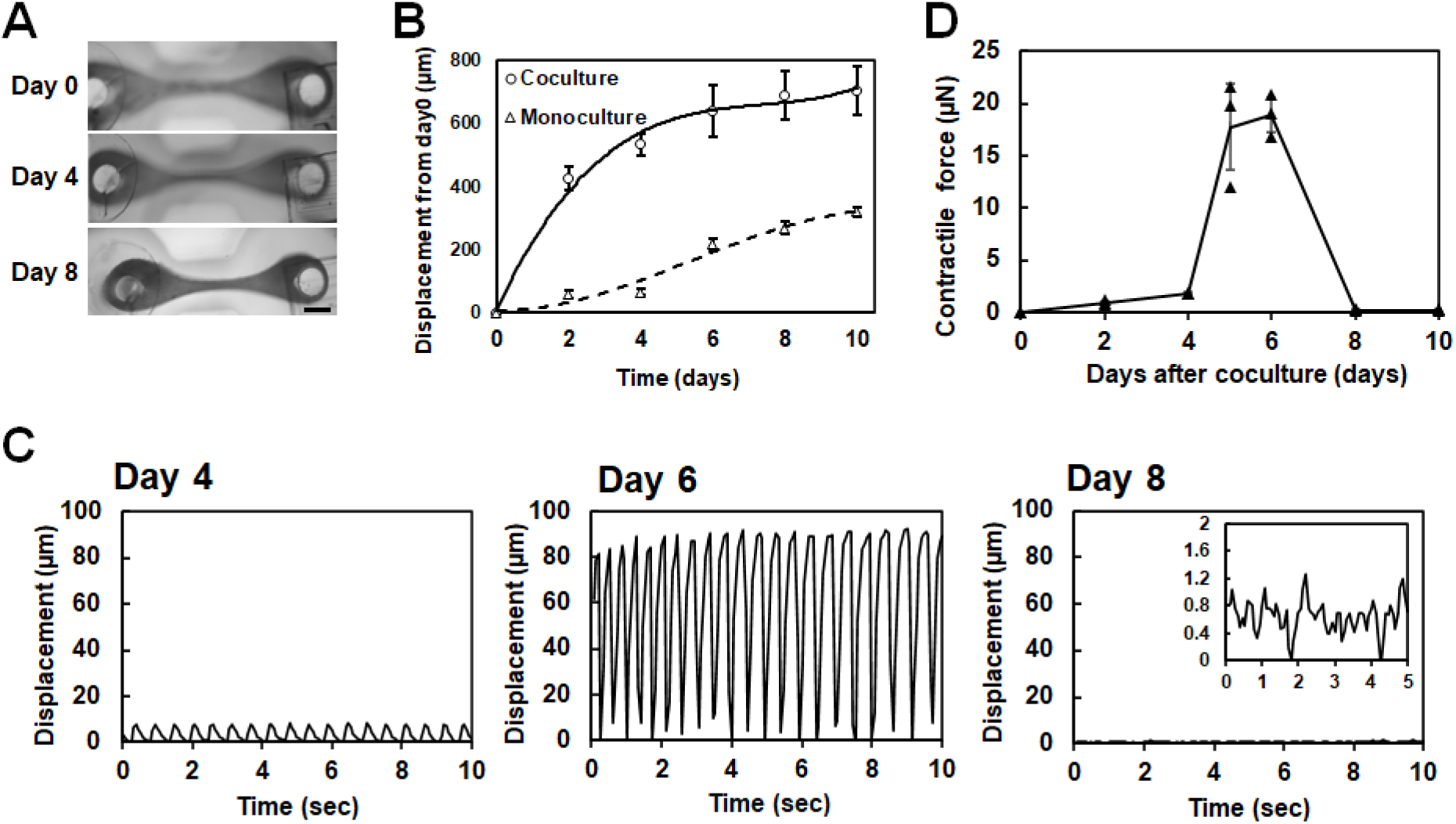
Coculture of MN spheroids and engineered SkM tissues on the device. A) Representative images of the SkM tissues on the device on day 0, 4, and 8. Scale bar; 500 μm. B) Displacement of the micropost by passive tension of the cocultured tissues. Open circle; with MN spheroids. Open triangle; without MN sheroids. C) Displacement of the micropost moved by the contraction of the SkM tissues which was generated in response to the spontaneous firing of the MNs on day 4, 6, and 8. D) Quantified contractile force of the SkM tissues cocultured with MNs on the devices.

During muscle development and regeneration *in vivo*, gap junctions that enable the propagation of electrical activity between cells are expressed transiently (36, 37). Furthermore, by *in vitro* experiments using 2D cultured rat primary myoblasts, Gorbe et al. reported that the permeability of gap junctions increased and decreased significantly in a few days after inducing differentiation (38). In this study, the contractile force of the cocultured tissues on the device reached a maximum value on day 6 and largely decreased on day 8. The periodicity was diminished on day 8 (Fig. 5C and D). Thus, we assume that the myotubes within the cocultured tissue on day 6 contracted synchronously in response to the spontaneous firing of MNs due to calcium ion propagation through the highly expressed gap junctions and, as the expression decreased, the myotubes contracted without synchronization, leading to the small and nonperiodical contraction of the tissues.

To evaluate whether the cocultured SkM tissues were innervated by MNs, we applied shear stress to the middle of the axons using the microchannel for axon stimulation (# in Fig. 2C) and observed the effects on the contraction of the tissues. As expected, the spontaneous contraction of the cocultured SkM tissues due to the spontaneous firing of MNs disappeared upon the application of shear stress (Fig. 6). This result shows that the myotubes in the SkM tissues on day 10 formed functional NMJs with the terminal of the axons from MN spheroids. This also indicates the possibility of using the developed device to model skeletal muscle atrophy after a nerve injury.

**Fig. 6.**
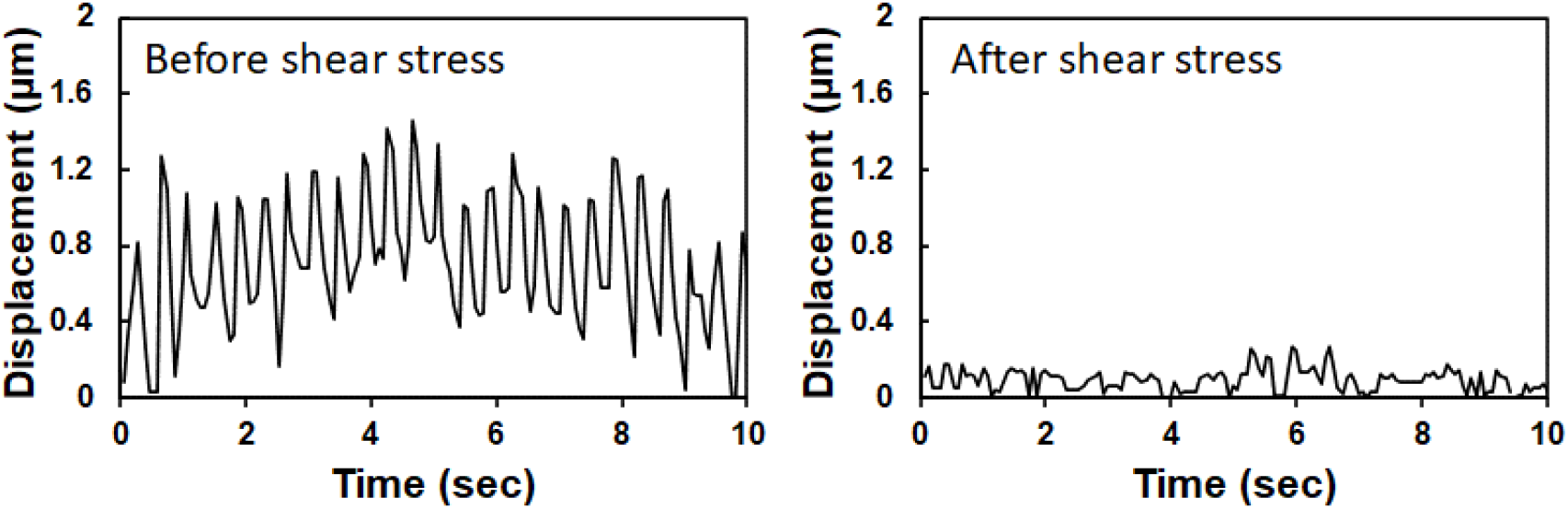
Application of the shear stress to the middle of the axon of the cocultured tissues on day 10. Representative results are displayed (n=3).

### 3.5 Functional assessments of the compartmentalized 3D neuromuscular tissue models on the devices

We evaluated the functionality of compartmentalized neuromuscular tissue models fabricated on the device. We added the excitatory neurotransmitter glutamate into the MN chamber to induce the firing of the MNs and measured the displacement of the micropost due to the SkM tissue contraction. We used the cocultured tissues on day 10. To reduce the spontaneous firing of MNs, the cocultured tissues were incubated at 25 °C for 10 min before adding glutamate.

As shown in Fig. 7A, the addition of glutamate into the MN chamber induced a lasting contraction, or tetanus, of the tissues. Subsequently, we added the sodium channel blocker, TTX, or antagonist to the nicotinic AChRs, Vb, into the MN chamber or SkM chamber, and evaluated their effects on the muscle contraction induced by glutamate (Fig. 7B and C). When TTX was added to the MN chamber, the muscle contraction did not occur, but muscle relaxation was observed (Fig. 7B-i). In contrast, when TTX was added to the SkM chamber, the muscle contraction was observed (Fig. 7B-ii). When Vb was added to the MN chamber, the muscle contraction was observed (Fig. 7C-i). In contrast, when Vb was added into the SkM chamber, the muscle contraction did not occur and muscle relaxation was observed (Fig. 7C-ii). The difference observed for TTX and Vb (Fig. 7B and C) could be explained by the inhibitory mechanisms of TTX and Vb. TTX blocks the voltage-gated sodium channels expressed in the MNs, while Vb binds to the nicotinic AChR expressed in the SkM cells. Thus, the compartmentalized neuromuscular tissue models fabricated on the device were functional. The results suggest that the tissue models could be used for phenotypic screening to evaluate the cellular type specific efficacy of drug candidates.

**Fig. 7.**
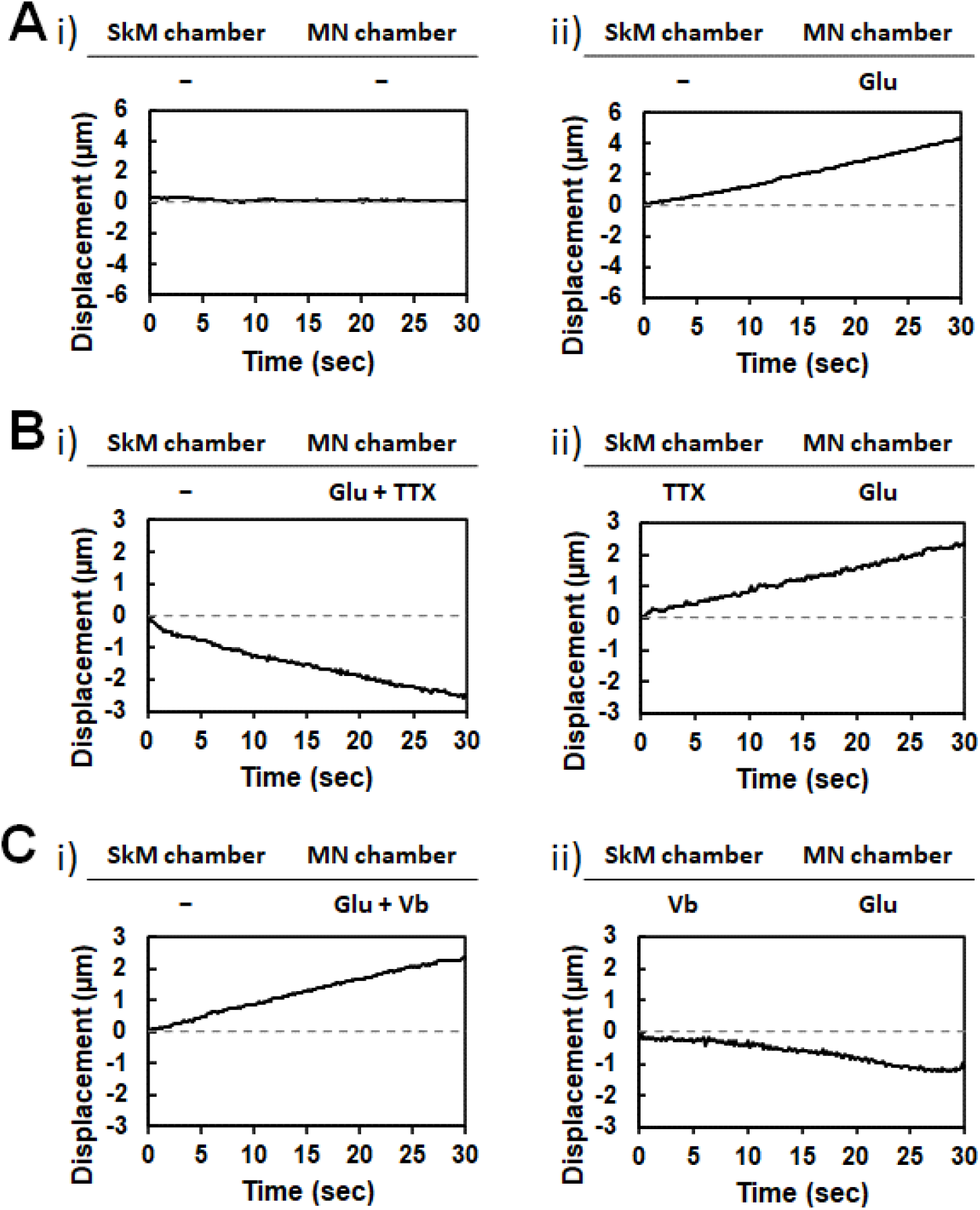
Assessment of the ftmctionality of the cocultured tissues on the devices. A) The displacement of the micropost after the addition of glutamate (Glu) into MN chamber. Representative results are displayed (n=3). B) The displacement of the micropost after the addition of TTX into SkM or MN chamber. Glutamate was added into the MN chamber before the addition of TTX into each chamber. Representative results are displayed (n=2). C) The displacement of the micropost after the additfon of Vb into SkM or MN chamber. Glutamate was added into the MN chamber before the addition of Vb into each chamber. Representative results are displayed (n=2).

As shown in Fig. 7A, the cocultured tissues on day 10 exhibited tetanic contraction by the addition of glutamate in the MN chamber. The previous 3D coculture models (8, 11, 14, 15) exhibited twitch contraction upon exposure to glutamate. Although the reason for the difference between the results of this study and previous studies is unclear, we believe that the maturation state of the SkM tissues in this study might be one of the reasons. Considering the precise compartmentalized nature of our device, the MNs and SkM tissues can be cultured in each specific culture medium, leading to the maturation of the SkM tissues. Furthermore, in this study, we used immortalized human myogenic cells, Hu5/KD3. In our previous study, we fabricated engineered Hu5/KD3 skeletal muscle tissues and showed that they generated a larger contractile force than that of the tissues engineered from other cell types, including C2C12, primary cells, and reprogrammed stem cells (28, 29). This result suggests that the maturation of Hu5/KD3 tissues was more advanced than those of C2C12 (8, 11, 14) or reprogrammed stem cells (15) used in other studies. Further experiments will be performed to understand the mechanisms underlying this phenomenon in the future.

## 4. Conclusion

In this study, we developed a microdevice for precisely compartmentalized coculturing of MNs and engineered SkM tissues. Our data suggest that the developed device has the following characteristics. First, it is possible to measure the force of engineered SkM tissues whose contraction is induced by MNs through NMJs. Second, it is possible to compartmentalize the cell bodies of MNs and SkM cells during a culture period. Third, it is possible to apply chemical/mechanical stimuli to the cell bodies of MNs and SkM cells and middle of the axons. Thus, the microdevice would be a useful tool in fundamental research and drug development for neuromuscular disorders.

## Acknowledgments

We thank Dr. Naohiro Hashimoto at the National Center for Geriatrics and Gerontology for providing Hu5/KD3 cells. This research was supported in part by Japan Society for the Promotion of Science KAKENHI Grant Numbers 26630429, JP16KK0126, JP18H01796, and JP20H04705. We would like to thank Editage for the English language editing.

---

**Supplementmy Fig. 1.**
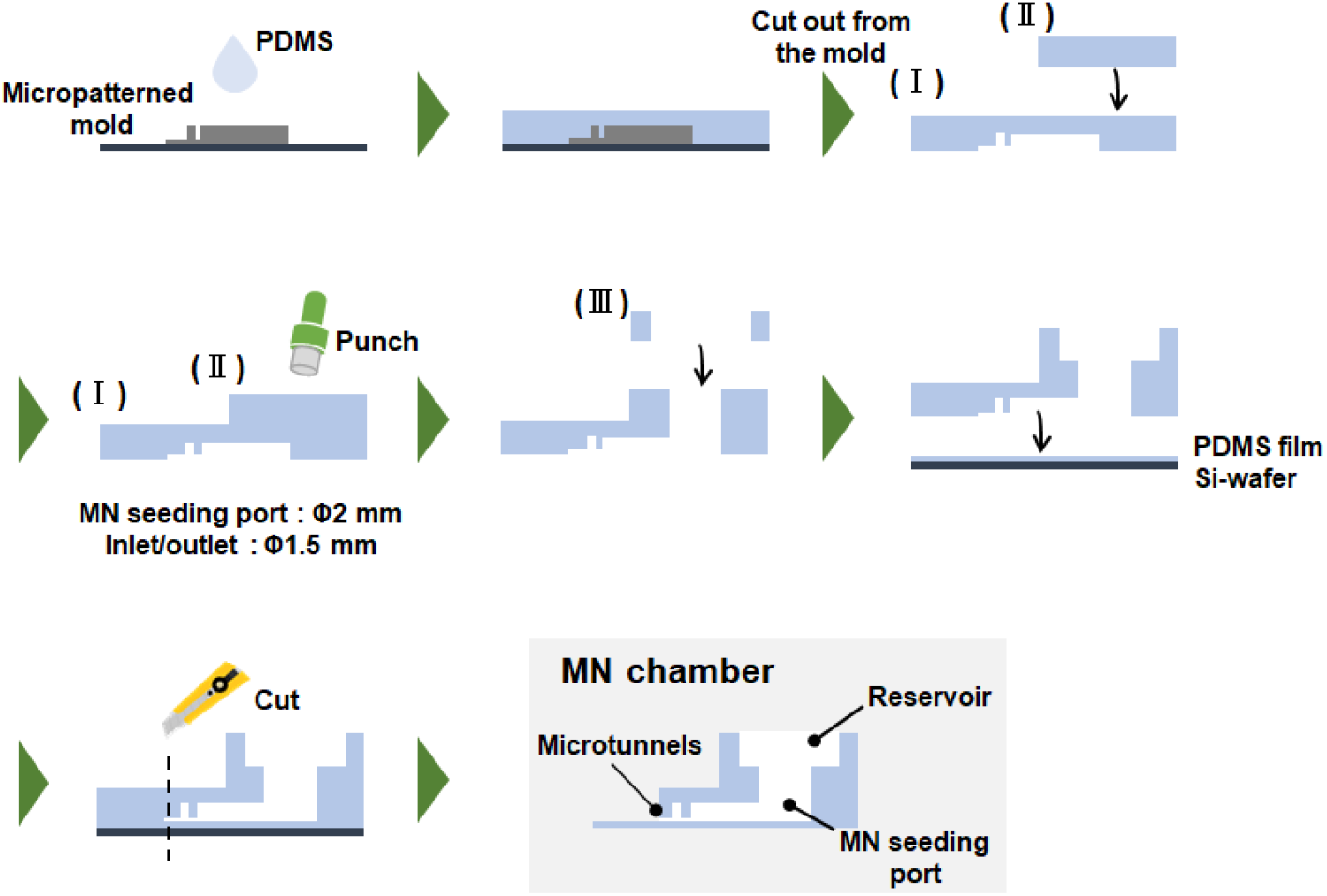
Fabrication process of the microdevice

**Supplementa1y Fig.2.**
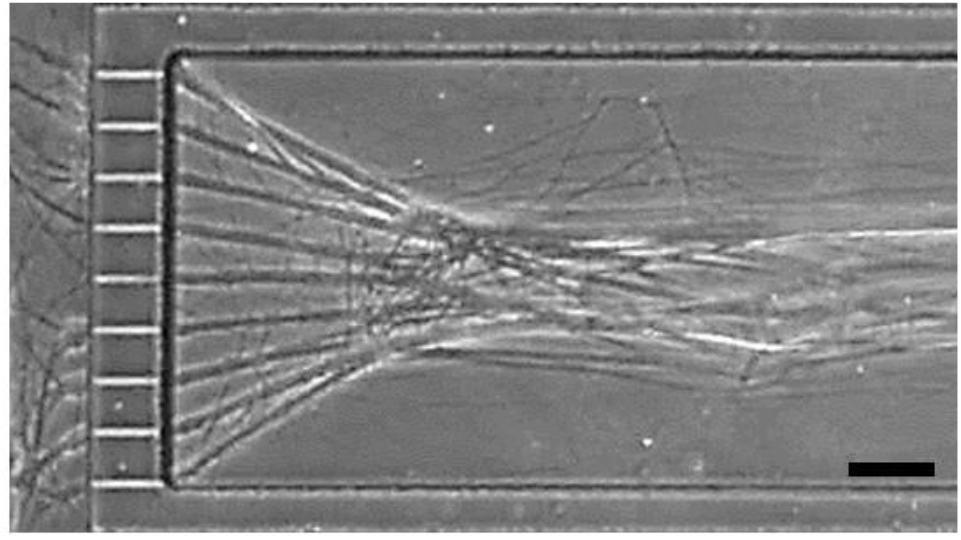
Axon elongation in the MN chmnber with 1 step 9 microtunnels for axons. Scale bar; 100 μm.

